# Structures of two LarA-like nickel-pincer nucleotide cofactor-utilizing enzymes with a single catalytic histidine residue

**DOI:** 10.1101/2025.08.19.671153

**Authors:** Santhosh Gatreddi, Sundharraman Subramanian, Dexin Sui, Tianqi Wang, Julian Urdiain-Arraiza, Benoit Desguin, Robert P. Hausinger, Kristin N. Parent, Jian Hu

## Abstract

The nickel pincer nucleotide (NPN) cofactor catalyzes the racemization/epimerization of α-hydroxy acids in enzymes of the LarA family. The established proton-coupled hydride transfer mechanism requires two catalytic histidine residues that alternately act as general acids and general bases. Notably, however, a fraction of LarA homologs (LarAHs) lack one of the active site histidine residues, replacing it with an asparaginyl side chain that cannot participate in acid/base catalysis. Here, we investigated two such LarAHs and solved their cryo-electron microscopic structures with and without loaded NPN cofactor, respectively. The structures revealed a consistent octameric assembly that is unprecedented in the LarA family and unveiled a new set of active site residues that likely recognize and process substrates differently from those of the well-studied LarAHs. Genomic context analysis suggested their potential involvement in carbohydrate metabolism. Together, these findings lay the groundwork for expanding the breadth of reactions and the range of mechanisms of LarA enzymes.

## Introduction

The organometallic nickel-pincer nucleotide (NPN) cofactor is composed of a nickel ion pincered by two sulfur and one carbon atoms from a pyridinium 3,5-dithiocarboxylate mononucleotide (P2TMN).^1,2^ The NPN cofactor, first discovered in the lactate racemase, LarA, from *Lactiplantibacillus plantarum* (LarA_*Lp*_),^3^ is biosynthesized by LarB,^4,5^ LarE,^6-8^ and LarC^9,10^ from nicotinic acid adenine dinucleotide,^11^ a precursor of nicotinamide adenine dinucleotide (NAD). The crystal structure of LarA_*Lp*_ reveals a covalent linkage of the NPN cofactor to a lysine residue via a thioamide bond, and shows the pincered nickel ion with a histidine residue completing the square-planar coordination.^3^ Situated over the NPN cofactor of LarA_*Lp*_ are two highly conserved histidine residues (His108 and His174) that function alternately as general acid and base during catalysis, according to the proposed proton-coupled hydride transfer (PCHT) mechanism (**Scheme 1**).^2,3,12^ Despite large sequence variation, LarA homologs (LarAHs) were thought to operate via the same PCHT mechanism, which has been supported by structural biology studies of representative LarA family members. For the family founding member LarA_*Lp*_, His108 and His174 bind sulfite, which is a strong competitive inhibitor and also an electron donor that can form a covalent S-C bond with the cofactor.^12,13^ A recent study of a LarAH from *Isosphaera pallida* reported the high-resolution structures in complex with three D-enantiomeric substrates, providing direct evidence supporting the catalytic role of the two histidines.^14^ Bioinformatic analysis has led to the identification of more than twenty LarA subfamilies in prokaryotic species, and the genomic context-guided biochemical study revealed diverse NPN cofactor-dependent racemization/epimerization reactions for various α-hydroxy acids.^15^ LarAHs from eukaryotic species were also recently reported,^16^ indicating the distribution of NPN cofactor-dependent enzymes across the three kingdoms of life. The greatly expanded substrate spectrum unveiled in that study underscores the importance of LarAs in the metabolism of α-hydroxy acids and carbohydrates.^16^

Notably, although most LarAHs possess two catalytic histidine residues that are essential for the PCHT mechanism, the second histidine residue (His174 in LarA_*Lp*_) in a small fraction of LarAHs is substituted by other amino acids, often an asparagine which cannot function as a general acid/base in catalysis. None of these LarAHs have been characterized, leaving unsolved questions concerning the structure-function relationship and catalytic mechanism of these putative enzymes. In this work, we report the cryo-electron microscopy (cryo-EM) structures of two such LarAHs in the NPN cofactor-free (apo) and NPN cofactor-loaded (holo) states, respectively, revealing an octameric state and a different active site from the other known LarAHs. Structural analysis, together with genomic context analysis, allowed us to propose their potential involvement in carbohydrate metabolism.

## Results and discussion

### Identification of a LarA subfamily that replaces a catalytic histidyl with an asparaginyl group

An early bioinformatic analysis reported three groups of LarAHs that either lack the catalytic histidine residue equivalent to His174 in LarA_*Lp*_ (Groups 14 and 15) or substitute it with an asparaginyl side chain (Group 17).^15^ Using the LarAH from *Blautia wexlerae* (LarA_*Bw*_) as seed, we conducted a BLAST search and identified hundreds of sequences that have an invariable asparagine residue at the position of His174 in LarA_*Lp*_. Sequence alignment of the selected LarAHs with identity as low as 29% revealed conserved residues in both N- and C-terminal domains (**Figure 1**), which are responsible for the binding of the NPN cofactor and substrate, respectively. Many residues conserved in this group of LarAHs, including the asparagine residue replacing the catalytic histidyl group, differ from those in other LarA subfamilies (**Figure S1**). Phylogenetic analysis showed that these LarAHs, represented by LarA_*Bw*_ and the corresponding protein from *Streptococcus plurextorum* (LarA_*Sp*_), form a distinct branch from other LarAHs with known substrates (**Figure S2**). Thus, these LarAHs likely form a unique subfamily that performs uncharacterized functions.

**Figure 1.**
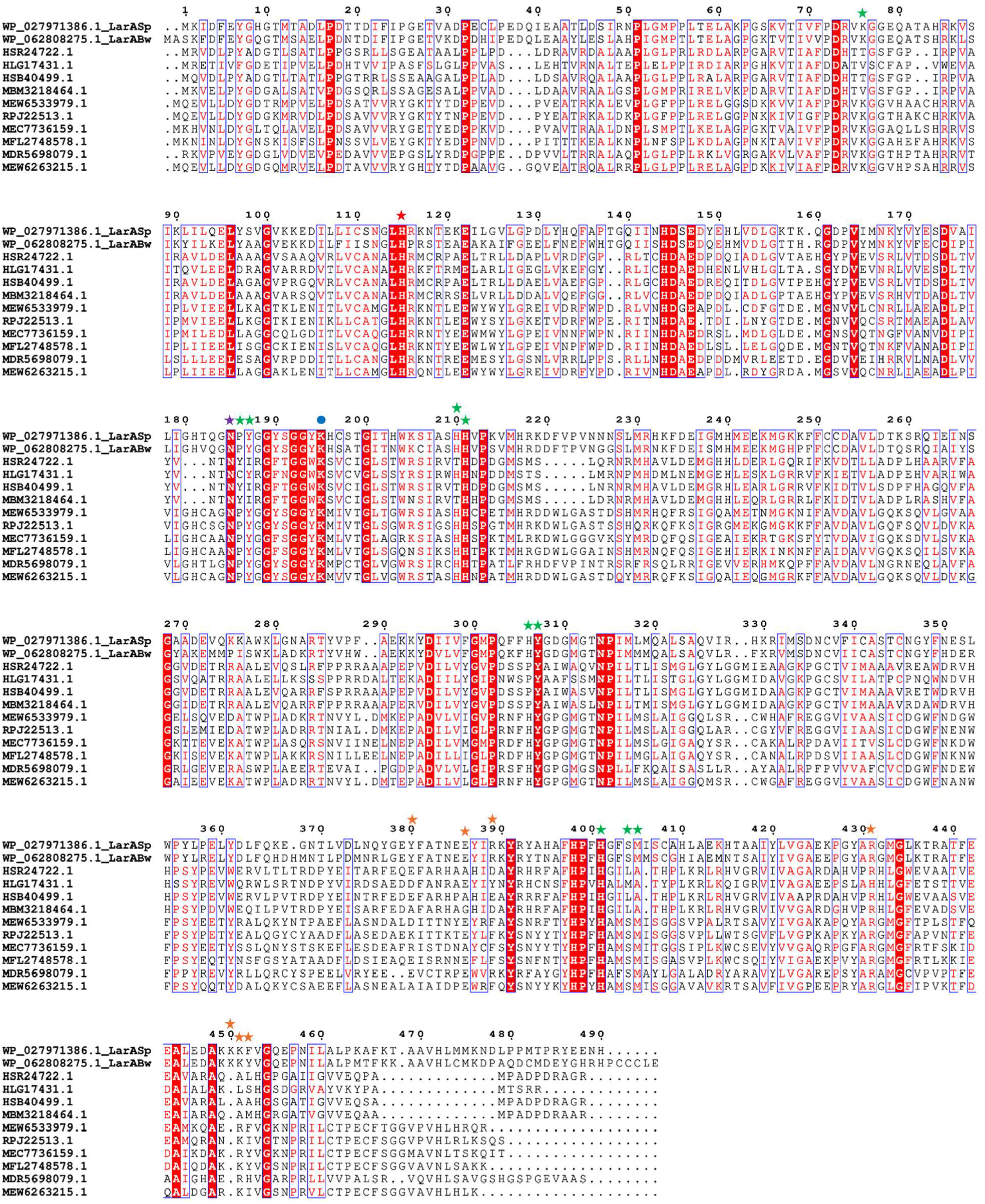
Multiple sequence alignment of LarAHs in a LarA subfamily with the second catalytic histidine residue substituted by an asparaginyl side chain. The catalytic histidine and the asparagine residues that are conserved in this subfamily are highlighted with red and purple stars, respectively. Other active site residues are indicated with green stars. Residues involved in the oligomerization are marked with orange stars and the lysine residue participating in covalent linkage with NPN is labeled with a blue dot. The LarAH protein IDs and their respective species are listed below. HSR24722.1: *Candidatus* Eisenbacteria; HLG17431.1: *Blastocatellia*; HSB40499.1: *Candidatus* Methylomirabilota; MBM3218464.1: *Candidatus* Rokubacteria; MEW6533979.1: *Thermodesulfobacteriota*; RPJ22513.1: *Desulfobacteraceae*: MEC7736159.1: *Candidatus neomarinimicrobiota*; MFL2748578.1: Unidentified eubacterium; MDR5698079.1: *Candidatus* Caldifonticola tengchongensis; MEW6263215.1: *Thermodesulfobacteriota*.

### Cryo-EM structure of LarA_Bw_ in the apo state

For structural study, the NPN cofactor is often incorporated *in vivo* by co-expression with the NPN synthesizing enzymes (LarB/C/E) in *L. lactis*.^3,12-14^ *In vivo* incorporation with coexpressed Lar proteins in *E. coli* or *in vitro* incorporation with biosynthesized NPN cofactor is an option for functional study but not recommended for structural study due to the relatively low incorporation efficiency.^11,15-17^ One protein that we characterized from this newly-identified LarA subfamily is a putative enzyme from a commensal human gut bacterium *B. wexlerae*. However, co-expression with the NPN-synthesizing enzymes in *L. lactis* produced only colorless protein, indicating that the purified LarA_*Bw*_ is in the apo state because otherwise the LarAH would be yellow or brown due to the NPN cofactor. While it is not uncommon that the NPN cofactor fails to be efficiently incorporated into a LarAH *in vivo*,^18^ this unsuccessful attempt limited our characterization of LarA_*Bw*_ to the apo form. For a better yield, The His_6_-tagged protein was overexpressed in *E. coli* and purified to homogeneity using immobilized metal affinity chromatography and size exclusion chromatography (SEC) (**Figure 2A**). Sodium dodecyl sulfate–polyacrylamide gel electrophoresis (SDS-PAGE) indicated a single band consistent with the theoretical molecular weight of 57.5 kDa. Based on the SEC elution profile and comparison to marker proteins, the purified LarA_*Bw*_ was in a large oligomeric state with an apparent molecular weight of ∼360 kDa.

**Figure 2.**
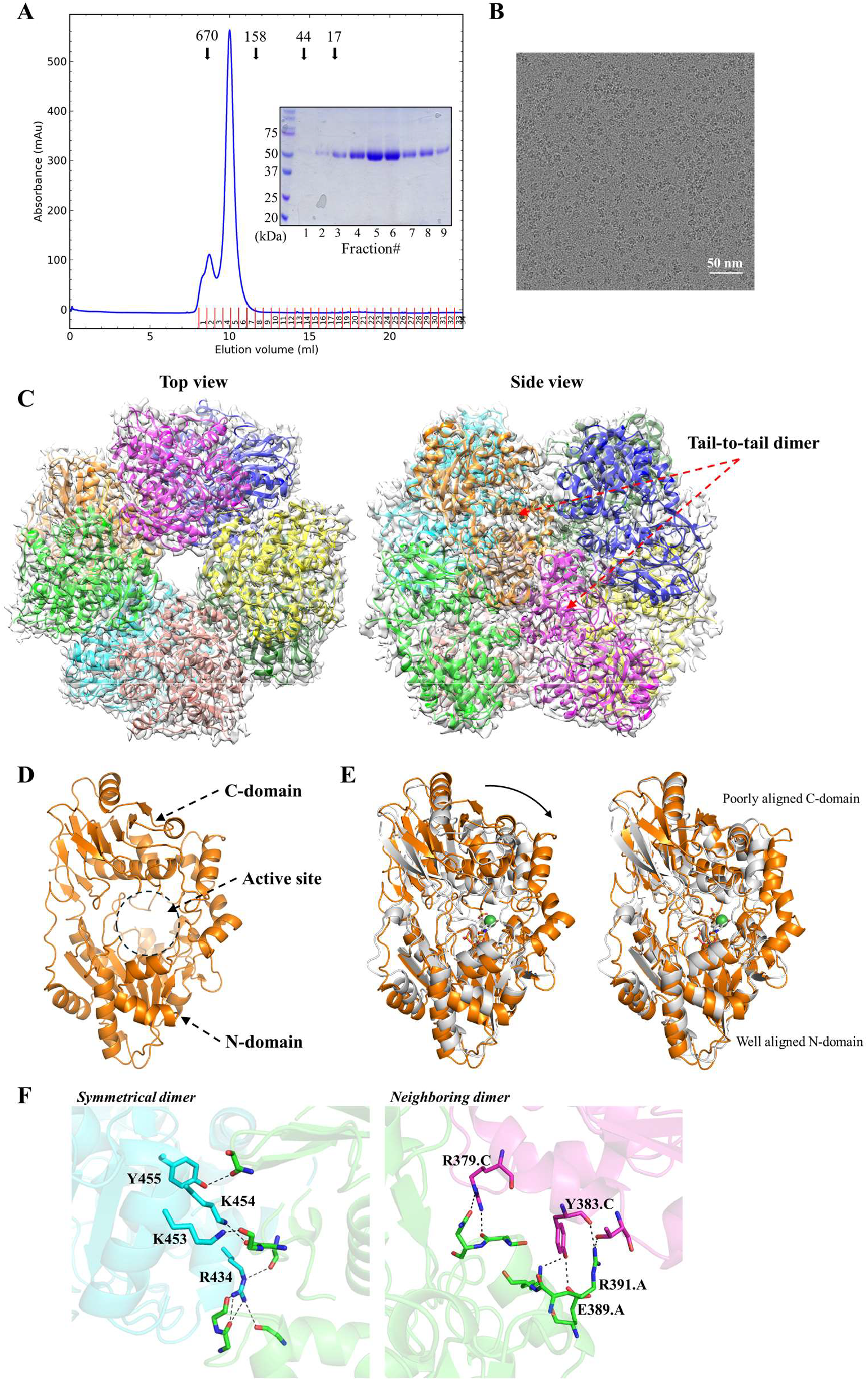
Cryo-EM structure of LarA_*Bw*_. (**A**) The SEC elution profile of LarA_*Bw*_. The protein standards, including thyroglobulin 670 kDa, gamma-globulin 158 kDa, ovalbumin 44 kDa, and myoglobulin 17 kDa, were eluted at the volumes indicated by the arrows. The SDS-PAGE profile of the peak fractions is shown in the inset. (**B**) A representative cryo-EM micrograph of LarA_*Bw*_. (**C**) Top view and side view of the octameric LarA_*Bw*_ in the density map. The basic structural unit is the tail-to-tail dimer as indicated in the side view. (**D**) Structure of the monomeric LarA_*Bw*_. The N- and C-domains of chain A, as well as the putative active site in between, are indicated. (**E**) Structural comparison of LarA_*Bw*_ and LarA_*Lp*_. *Left*: structural superposition of LarA_*Bw*_ (chain A, orange) and LarA_*Lp*_ (5HUQ, chain B in gray). The NPN cofactor in LarA_*Lp*_ is shown in stick mode with the nickel depicted as a green sphere. The curved arrow indicates a more open conformation of LarA_*Bw*_ in the apo state. *Right*: structural superposition of separated N- and C-domains, revealing an overall good structural alignment for the N-domains but poor alignment for the C-domains. (**F**) Zoomed-in view of the interfaces stabilizing the octamer. *Left*: polar contacts stabilizing the tail-to-tail dimerization. *Right*: polar interactions between the neighboring subunits in a tetramer. Note that the conserved R434 is involved in multiple hydrogen bonds. Only the interfacial residues that use their side chains to form polar interactions are labeled and shown in stick mode. The polar interactions are indicated by dashed lines.

The oligomeric LarA_*Bw*_ was not crystallizable, but its structure was readily solved by cryo-EM with a C4 symmetry at a resolution of 3.15 Å, revealing an octameric assembly (**Figures 2B&2C**). Like other LarAHs, LarA_*Bw*_ is a two-domain protein with the putative active site located between the N- and C-terminal domains (**Figure 2D**). The Cα root mean square deviation (RMSD) of the monomeric LarA_*Bw*_ with LarA_*Lp*_ (PDB 5HUQ, chain B) is 4.2 Å, indicative of a significant structural difference (**Figure 2E**). Further analysis of the individual domains showed that the N-terminal domains of LarA_*Bw*_ and LarA_*Lp*_ could be reasonably aligned with a Cα RMSD of 1.25 Å, whereas the large portion of the C-terminal domains cannot be aligned despite the similar arrangement of the secondary structure elements. The large RMSD of 4.2 Å for the full-length proteins can also be attributed to the different domain orientations – LarA_*Bw*_ adopts a conformation with the active site more open to the solvent than LarA_*Lp*_. The detailed comparison of the active sites will be elaborated in a later section.

The basic structure unit of the octamer is a symmetrical dimer in which the C-terminal domain of each protomer contacts through extensive polar interactions (**Figure 2F**, left panel), burying 1076 Å^2^ surface area. Most of the polar interactions are mediated by the conserved Arg434 (**Figure 1**), which uses its guanidinium side chain to form hydrogen bonds with multiple backbone carbonyl oxygen atoms from the other protomer. Several other polar residues (Lys453, Lys454, Tyr455) are also involved in dimerization, but they are variable in this subfamily (**Figure 1**). In contrast, the interactions between the neighboring dimers are mediated by nonconserved residues with only 300-400 Å^2^ of surface area buried at the interface (**Figure 2F**, right panel), consistent with the notion that the octamer forms through the tetramerization of the dimers. Indeed, the RMSD values of the four dimers are smaller than 0.1 Å. A square-shaped cavity with a side length of ∼40 Å is found within the octamer with an entrance size of ∼16 Å at the centers of the top and bottom tetramers (**Figure S3**). As the surface or the entrance of the cavity is not lined with conserved residues, it is unlikely that the cavity is related to the function of LarA_*Bw*_.

### Production of LarA_Sp_ in the holo state

To generate a LarAH in this subfamily in the holo state, we cloned the gene for a homolog of LarA_*Bw*_ from *S. plurextorum*, a pathogen of swine, which shares 72% identical residues with LarA_*Bw*_, and we co-expressed the C-terminal strep-tagged protein with LarB/C/D/E in *L. lactis*. The purified LarA_*Sp*_ was eluted from SEC as a large oligomerwith nearly the same apparent molecular weight as the octameric LarA_*Bw*_ (**Figure 3A**). Importantly, the purified protein exhibited a yellow-brown color, which is consistent with the holo state of LarAHs. The UV-Vis spectrum displayed weak absorptions at 340 nm and 550 nm, and a stronger absorption at 420 nm (**Figure 3B**). The absorptions at 420 nm and 550 nm are characteristic for the NPN cofactor-loaded LarAHs, whereas the absorption at 340 nm implies that a fraction of the purified protein may contain the reduced NPN cofactor, which was observed only when lactate (substrate) or NaBH_4_ (hydride donor) was added to LarA_*Lp*_.^12,13^ These data suggest that the purified LarA_*Sp*_ may be bound with a ligand that perturbs the redox state of the NPN cofactor.

**Figure 3.**
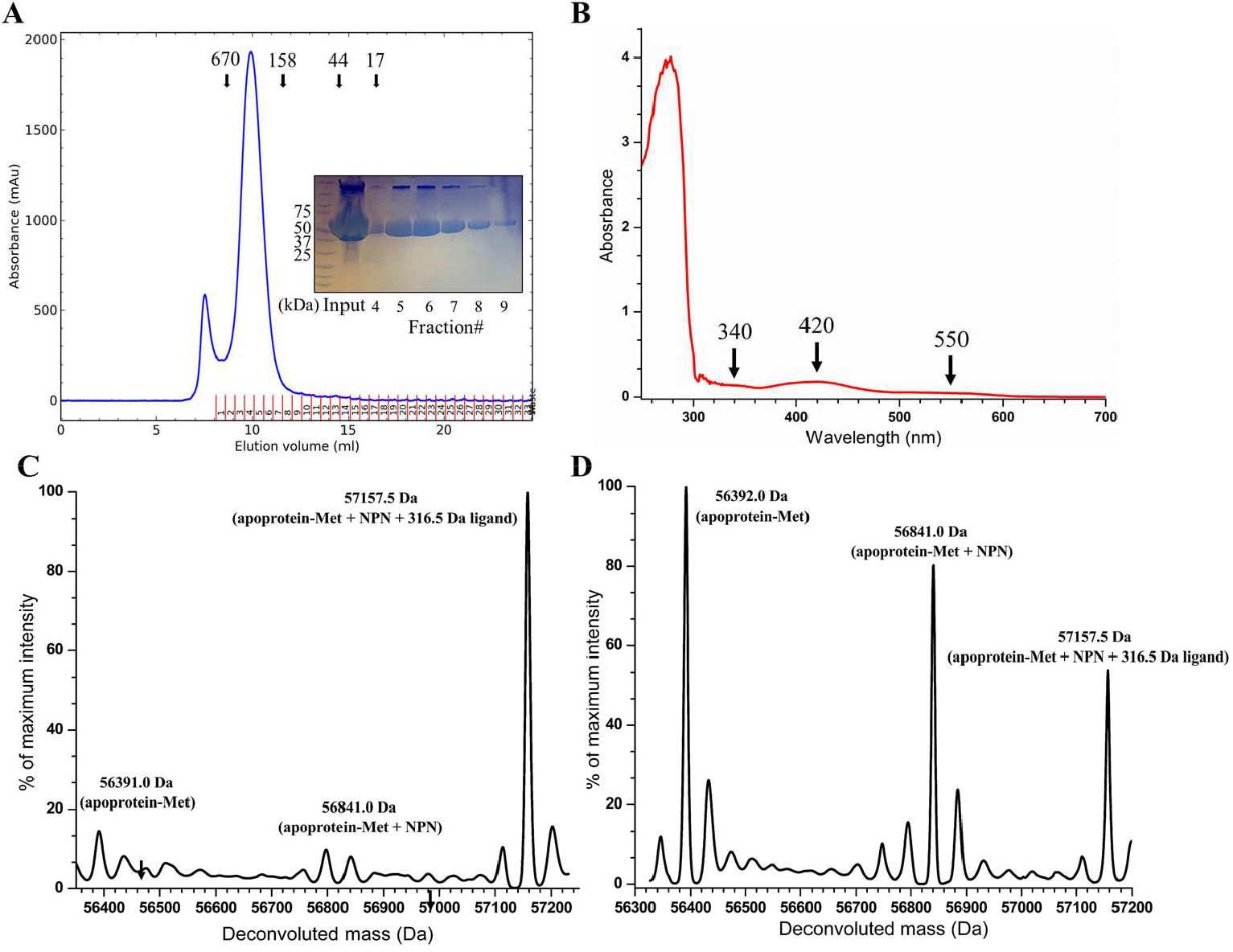
Production of LarA_*Sp*_ in the holo state. (**A**) The SEC elution profile of LarA_*Sp*_. The protein standards were eluted at the volumes indicated by the arrows. The SDS-PAGE profile of the peak fractions is shown in the inset. (**B**) UV-Vis spectrum of the purified LarA_*Sp*_. The major absorption features are indicated with arrows. (**C**) and (**D**) ESI-MS analysis of purified LarA_*Sp*_. The fresh sample is shown in (C) and an aged one is shown in (D).

Unexpectedly, electrospray ionization mass spectrometry (ESI-MS) analysis of the purified LarA_*Sp*_ revealed a major species with a molecule weight of 57,157.5 Da (**Figure 3C**), which is 316.5 Da greater than the sum of the protein in the apo state (assuming loss of the N-terminal methionine residue, 56,392.0 Da) and the NPN cofactor (449.0 Da). Of interest, analysis of a sample stored at the room temperature for one week showed that the major species degraded to the apo state protein and the holo state protein (i.e., the NPN cofactor-loaded state) (**Figure 3D**), suggesting that a covalently bound ligand with a molecular weight of 316.5 Da may be slowly released over time. Because the ESI-MS data demonstrated the holo state of the purified LarA_*Sp*_, we were encouraged to move forward with structural characterization.

### Structure determination of LarA_Sp_ in the holo state

The large molecular weight and C4 symmetry of LarA_*Sp*_ allowed us to solve the cryo-EM structure of LarA_*Sp*_ at a resolution of 2.2 Å (**Figures 4A&4B**). Like LarA_*Bw*_, LarA_*Sp*_ also assembles as an octamer that is a tetrameric dimer, and the monomeric protein has a putative active site between the N- and C-domains (**Figure 4C**). Similar to LarA_*Bw*_, a larger surface area (1130 Å^2^) is buried at the interface of two C-terminal domains than those between the neighboring monomers within the top or bottom tetramer (300-400 Å^2^), supporting the notion that the basic unit of the octamer is a dimer. Arg431 (equivalent to Arg434 in LarA_*Bw*_) forms multiple hydrogen bonds with the same set of residues from the other protomer as in LarA_*Bw*_ at the dimerization interface, suggesting that the tail-to-tail dimer is common in this LarA subfamily whereas octamerization may not be so prevalent since the residues at the interface between the neighboring dimers are not conserved. The octameric LarA_*Bw*_ and LarA_*Sp*_ are highly superimposable with a Cα RMSD of 2.0 Å (**Figure 4D**, left panel), and the tail-to-tail dimers and monomers from both structures are aligned with the Cα RMSD values of 1.6 Å and 0.94 Å, respectively. Therefore, the loading of the NPN cofactor does not change the oligomerization or the domain orientation in each monomer, except that the loop containing the asparagine residue of interest moves toward the NPN cofactor in the holo state (**Figure 4D**, right panel).

**Figure 4.**
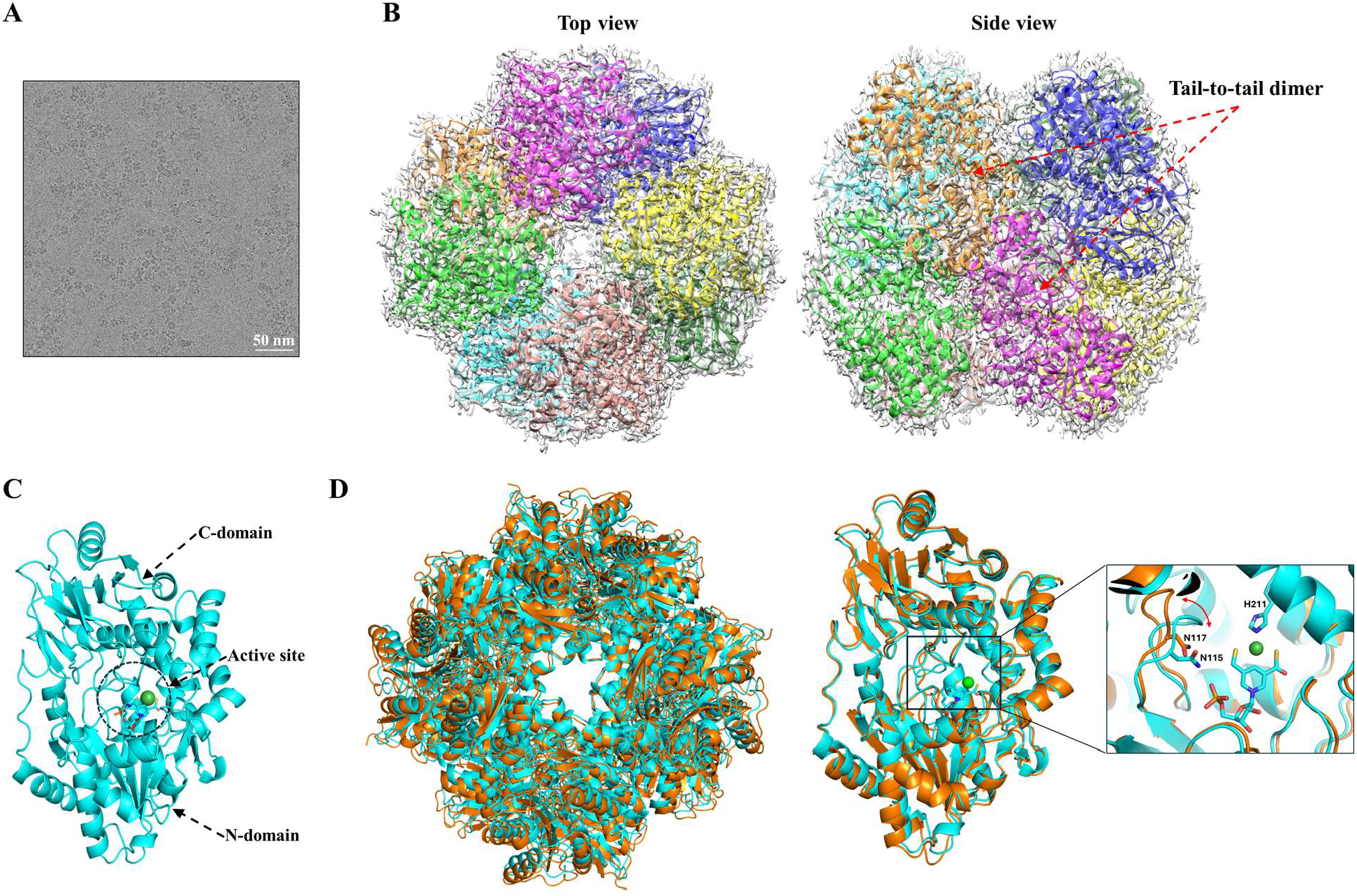
Cryo-EM structure of LarA_*Sp*_. (**A**) A representative cryo-EM micrograph of LarA_*Sp*_. (**B**) Top view and side view of the octameric LarA_*Sp*_. (**C**) Structure of the monomeric LarA_*Sp*_. The N- and C-domains of chain A, as well as the putative active site in between, are indicated. (**D**) Structural comparison of LarA_*Sp*_ (cyan) and LarA_*Bw*_ (orange). *Left*: superposition of octamers. *Right*: superposition of monomers. Insert: zoomed-in view of the active site, highlighting the displacement of the loop containing the histidine-substituting asparagine residue with the red curve arrow. The NPN cofactor and the asparagine residues are shown in stick mode and the nickel ion is depicted as a green sphere.

At the active site, a continuous density matches the NPN cofactor (**Figure 5A**). As this density is connected to the side chain of Lys195, the NPN cofactor forms a covalent linkage through a thioamide bond to the enzyme. Like in LarA_*Lp*_ and LarA_*Ip*_, the nickel atom in the NPN cofactor is coordinated to the C4 atom of the pyridinium ring, two sulfur atoms, and the imidazole side chain of His211 in a square-planar coordination (**Figure 5B**). Over the NPN cofactor, we noticed electron density in close proximity to the conserved His115, His306, Tyr307, and His401, occupying a position corresponding to the substrate binding sites of other LarAHs (**Figure S4**). This density also extends toward the entrance of the active site. Considering the 340 nm absorption in the UV-Vis spectrum (**Figure 4C**) and the 316.5 Da extra mass (**Figures 4D&4E**), we speculate that an unidentified ligand is covalently bound at the substrate binding site. However, we were not successful at determining the identity of this ligand due to its low occupancy, as suggested by the poor density map.

**Figure 5.**
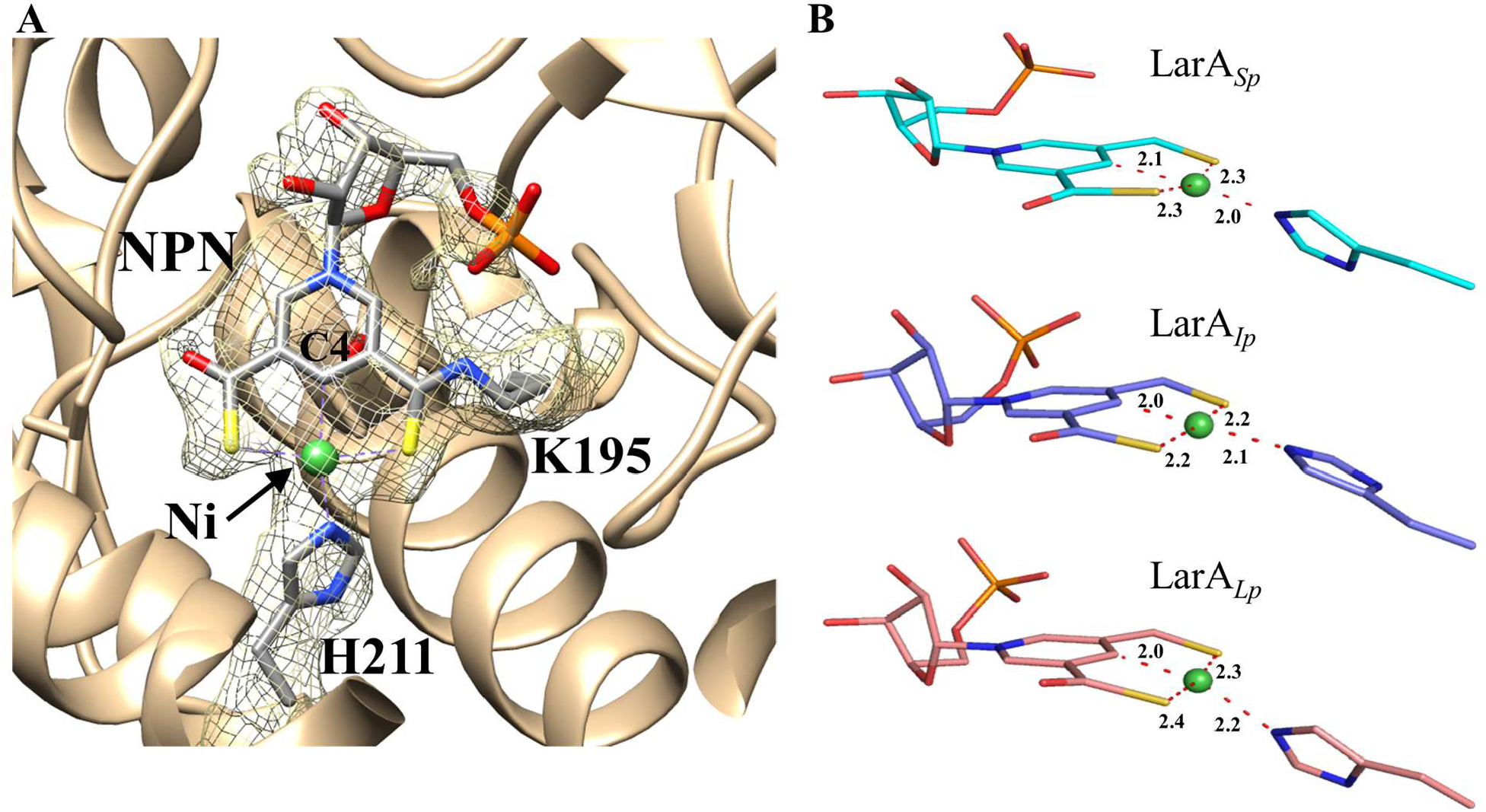
Nickel coordination in the NPN cofactor of LarA_*Sp*_. (**A**) Electron density of the NPN cofactor (chain D). The density of the cofactor is connected to that of Lys195, indicating a thioamide bond. His211 is the fourth ligand that completes the distorted square-planar coordination of the nickel ion. (**B**) Comparison of Ni coordination in LarA_*Sp*_, LarA_*Ip*_ (PDB 9EIA, chain A), and LarA_*Lp*_ (PDB 6C1W, chain B). The distances between Ni and coordinating atoms are labeled in angstrom.

In parallel to the cryo-EM study, we also crystallized LarA_*Sp*_ and solved the structure at 3.1 Å. Crystal packing analysis showed that LarA_*Sp*_ forms the same octamer as revealed in the cryo-EM structure (**Figure S5**). The basic tail-to-tail dimers from the two octamers are highly superimposable with a Cα RMSD of 1.1 Å. However, due to the low resolution, the density in the x-ray structure was too poor to model the NPN cofactor or the unidentified ligand.

### Comparison of the active sites

The active sites of LarA_*Sp*_ and LarA_*Bw*_ were compared with that of LarA_*Ip*_ (**Figure 6**). Only a few residues in the active sites are conserved among these LarAHs. They include a catalytic histidyl side chain (His115, His117, and His107 in LarA_*Sp*_, LarA_*Bw*_, and LarA_*Ip*_, respectively), the histidine residue for Ni coordination (His221, disordered His223, and His199), the lysyl group for the covalent linkage with the NPN cofactor (Lys195, Lys197, and Lys183), and the aspartate residue associating with the ribose moiety of the cofactor (Asp73, Asp75, and Asp70). In addition, Arg73 for the binding of the NPN phosphate group and the carboxylate group of D-lactate in LarA_*Ip*_ is conservatively substituted with Lys76 in LarA_*Sp*_ and Lys78 in LarA_*Bw*_, respectively.

**Figure 6.**
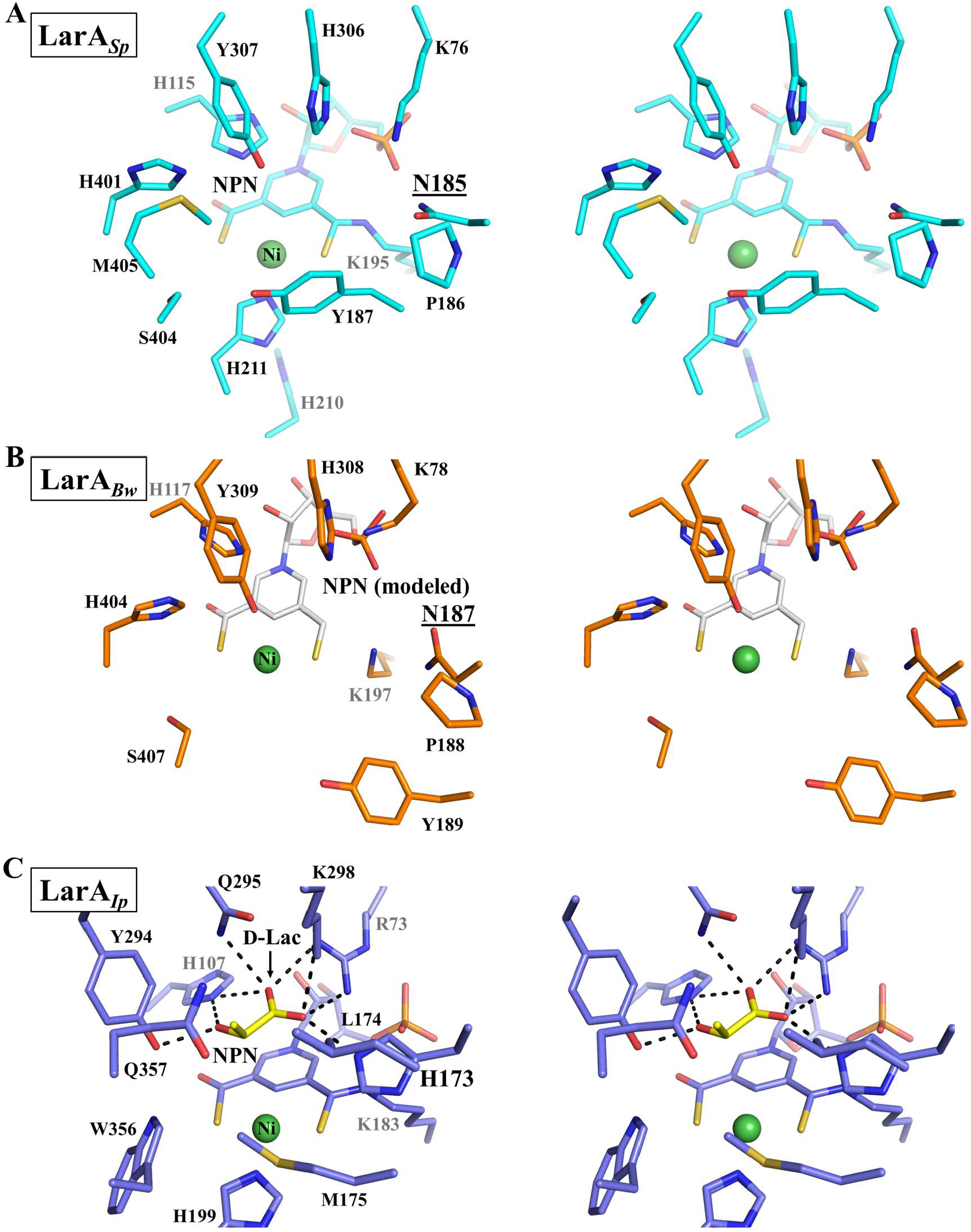
Stereo views of the active sites of (A) LarA_*Sp*_ (cyan), (B) LarA_*Bw*_ (orange), and (C) LarA_*Ip*_ (blue). LarA_*Ip*_ (PDB 9EIA, chain A) is bound to an authentic substrate, D-lactate (D-Lac, yellow). Asn185 in LarA_*Sp*_ and Asn187 in LarA_*Bw*_, which substitute for the catalytic histidine residue in LarA_*Ip*_ (His173), are underlined. An NPN cofactor (white) is modeled in the structure of LarA_*Bw*_ for better comparison. Some residues for nickel or substrate binding in LarA_*Bw*_ were missing due to lack of electron density.

Besides these residues that are primarily involved in the binding of the NPN cofactor, other active site residues in LarA_*Ip*_ are not identical to or conservatively substituted with those in the new subfamily members. The most notable difference is that the second catalytic histidyl group (His173 in LarA_*Ip*_) is substituted by Asn195 in LarA_*Sp*_ and Asn197 in LarA_*Bw*_. According to the PCHT mechanism (**Scheme 1**), His107 and His173 in LarA_*Ip*_ alternately function as a general base to deprotonate 2-OH of D/L-enantiomeric α-hydroxyacids and a general acid to protonate the pyruvate intermediate to complete racemization. Since the asparagine residue is absolutely conserved in the newly identified LarA subfamily (**Figure 1**) and topologically equivalent to the catalytic His173 in LarA_*Ip*_ according to the structural comparison, its inability to catalyze an acid/base reaction rules out the possibility of functional equivalence to His173 in catalysis. Given that His107 and His173 deprotonate D- and L-enantiomers respectively, the substitution of the L-enantiomer specific His173 with an asparagine residue suggest that LarA_*Sp*_ and LarA_*Bw*_ may only process D-enantiomer substrates and thus function as a unidirectional racemase/epimerase. Although uncommon, atypical racemases that process only one enantiomer have been reported. For instance, the cofactor independent aspartate/glutamate racemases usually use two cysteine residues to alternately act as a general base to deprotonate the chiral Cα to achieve racemization.^19,20^ Remarkably, some aspartate/glutamate racemases have only one cysteine residue with the other replaced by a non-cysteine residue. Biochemical and structural studies of such an enzyme from *E. coli* showed that it only catalyzes the conversion from an L-amino acid to a D-amino acid but cannot catalyze the D-to-L conversion, which has been attributed to the lack of the second catalytic cysteine residue.^21^ In addition to the lack of the catalytic histidine residue, many other active site residues of LarA_*Sp*_ and LarA_*Bw*_ are different from those in LarA_*Ip*_ (**Figure 6**), strongly suggesting that these putative enzymes may process chemicals that are very different from the known substrates of LarAs. Consistently, we were unable to detect any racemization activity of LarA_*Sp*_ for D-/L-lactate (data not shown), although it has been shown that many LarA homologs are highly promiscuous.^14,16^

### Potential involvement in carbohydrate metabolism

Genomic context analysis can facilitate identifying substrates of enzymes,^22^ including LarAHs.^15^ In this work, we examined the contexts of the genes encoding LarA_*Sp*_ and LarA_*Bw*_, respectively. Of interest, both *larA* genes have gene neighbors that potentially encode enzymes involved in glycolysis, including glucose-6-phosphate isomerase, fructose-6-phosphate-1-phosphotransferase and triose phosphate isomerase in *S. plurextorum*, and fructose-1,6-bisphosphatase, triose phosphate isomerase, fructose-bisphosphate aldolase, glucosamine-6-phosphate isomerase, and aldolase-1-epimerase in *B. wexlerae* (**Figure S6**). Several genes encoding subfamily members that share sequence identity as low as 37% (**Figure 1**) are also located next to the genes encoding possible sugar ABC transporters and enzymes involved in carbohydrate metabolism, allowing us to speculate that these LarAHs are potential carbohydrate-processing enzymes. Based on the density map and the MS data, we modeled a linear phosphorylated aldose with a formula of C_9_H_19_O_10_P (316.2 Da for the deprotonated form) to fit the density (**Figure S7**). Although this model is highly speculative, it suggests the size and shape of this unidentified ligand.

## Conclusion

In this study, we characterized two members of a newly identified LarA subfamily that lacks one of the catalytic histidine residues conserved in other LarAHs. Using cryo-EM, we determined their structures in the apo and holo states at atomic resolutions. From the structural data, combined with MS and spectroscopic analysis, as well as genomic context analysis, our findings revealed (1) a unique octameric assembly which is unprecedented among other known LarAHs; (2) a distinct set of active site residues, including the asparagine side chain substitution for the missing histidyl group, suggesting recognition and processing of non-canonical LarA substrates; and (3) genomic and structural evidence hypothetically linking these LarAHs to carbohydrate metabolism. This work provides a foundation for exploring the biological functions and catalytic mechanisms of these putative enzymes potentially involved in carbohydrate metabolism.

## Methods

### Genes and constructs

To express LarA_*Bw*_ in *L. lactis*, a PciI restriction enzyme site was first introduced into the pGIR112 vector^3^ using primers PciI_F and Pcil_R, and then the PCR product of the gene encoding LarA_*Bw*_ amplified using primers LarAH13_F and LarAH13_R was inserted into the plasmid via PciI and NheI, generating the plasmid pGIR112_LarA_*Bw*_. The gene encoding LarA_*Sp*_ was PCR amplified using primers LarAH37_A and LarAH37_B2, digested by XbaI and PciI, and inserted into the pGIR210 vector^15^ that had been digested with NheI and PciI, generating the plasmid pGIR213 for coexpression of LarA_*Sp*_ with LarB/C/E in *L. lactis*. The plasmids and primers used in this study are listed in **Table S1**.

### Protein expression and purification

Expression and purification of LarA_*Bw*_. The *E. coli* strain BL21-Gold(DE3) bearing the pET23b plasmid expressing LarA_*Bw*_^23^ was grown overnight at 37 °C and 220 rpm in lysogeny broth (LB) containing 100 µg/ml of ampicillin. Overnight grown culture was diluted to 1% with LB broth supplemented with 100 µg/ml of ampicillin and allowed to grow at 37 °C with shaking at 220 rpm. When the optical density at 600 nm (OD_600_) reached 0.35, the culture was cold-shocked by incubating on ice for 20 min and then 0.1 mM IPTG was added, followed by shaking at 16 °C for 24 h. Cells were harvested by centrifugation at 6,000 rpm for 10 min. Pelleted cells were resuspended in a buffer containing 20 mM Tris, pH 8.0, 300 mM NaCl, and 5% glycerol. Cells was lysed by sonication and the supernatant was collected after centrifugation at 20,000 *g* and 4 °C for 35 min. His_6_-tagged LarA_*Bw*_ in the supernatant was sequentially purified using the Ni-NTA resin and then a Superdex 200 increase 10/300 GL column equilibrated with a buffer containing 10 mM HEPES, pH 7.8, and 150 mM NaCl. The peak fractions from SEC were pooled for cryo-EM grid preparation.

Expression and purification of LarA_*Sp*_. The *L. lactis* cells transformed with the pGIR213 plasmid were grown without shaking at 30 °C overnight in M17 media supplemented with 0.5% glucose and 7.5 µg/ml of chloramphenicol. The overnight grown culture was diluted to 1% with M17 medium containing 0.5% glucose and 7.5 µg/ml of chloramphenicol and grown at 30 °C with gentle shaking at 220 rpm until the OD_600_ reached 0.4-0.5. Protein expression was induced by adding 5 µg of Nisin A per liter culture for 4 h, after which the culture was stationed overnight at 4 °C before cell harvest by centrifugation at 4,000 *g* for 20 min. Pelleted cells were resuspended in a buffer containing 100 mM Tris, pH 7.5, 150 mM NaCl, 2.5 µg/ml of DNase I, and 15 µg/ml of lysozyme. Cell lysis was performed using a French Press at 15,000 psi by passing the cell suspension twice. Clarified supernatant was obtained after centrifugation at 18,000 *g* and 4 °C for one h and applied to Strep-Tactin®XT 4Flow® high capacity resin (IBA Lifesciences) pre-equilibrated with a buffer containing 100 mM Tris, pH 7.5, and 150 mM NaCl. Resin was washed with 100 mM Tris, pH 7.5, and 150 mM NaCl, and strep-tagged LarA_*Sp*_ was eluted with the wash buffer supplemented with 50 mM biotin. The elution fractions containing the protein of interest were concentrated using an Amicon Ultra centrifugal filter and then applied to a Superdex 200 increase 10/300 GL column. The peak fractions from SEC were pooled for cryo-EM grid preparation and crystallization.

### UV-Visible spectroscopy

The UV-Visible spectrum (250-700 nm) of purified LarA_*Sp*_ (23 mg/ml) was recorded on a Shimadzu UV-2600 spectrophotometer (Kyoto, Japan) with a 10 mm path length and a 2 nm slit width for two accumulations.

### ESI-MS analysis

Purified LarAH_*Sp*_ in 50 mM Tris, pH 7.5, and 125 mM NaCl was analyzed by ESI-MS. Data were collected in positive ion mode on Xevo G2-XS QTof (Waters) equipped with Thermo Hypersil Gold CN guard desalting column. 10 μl of protein sample was injected at a flow rate of 0.1 ml/min. The mobile phases were 0.1% formic acid in water (solvent A) and acetonitrile (solvent B) initially mixed at a 98%:2% ratio, and then solvent B was gradually increased to 75%. The MaxEnt1 algorithm was used to generate the molecular mass spectra.

### Cryo-EM grid preparation, data collection, and processing

Quantifoil R2/2 UT 200 mesh copper grids (for LarA_*Bw*_) or Quantifoil R1.2/1.3 200 mesh copper grids (for LarA_*Sp*_) were treated using a Pelco easiGlow™ glow discharge for 45 s. 5 µl of the sample at 2 mg/ml (for LarA_*Bw*_) or 3.5 µl of the sample at 1.5 mg/ml (for LarA_*Sp*_) was added to the grid. The grids were then blotted using a Vitrobot Mark IV system before being plunged into liquid ethane.

For LarA_*Bw*_, single particle cryo-EM data were collected at the Cryo-EM facility of Michigan State University using a Talos Arctica equipped with a Falcon 3 direct electron detector, operating at 200 keV. A total of 1,539 micrographs were collected at ×120,000 nominal magnification (0.872 Å/pixel) over 44.60 s for a total dose of 32.27 e^−^/Å^2^. For LarA_*Sp*_, single particle cryo-EM data were collected using the Talos Arctica equipped with a Falcon 4i direct electron detector, operating at 200 keV with a post-column Selectris energy filter (10-eV slit width). A total of 4,657 micrographs were collected at ×130,000 nominal magnification (0.886 Å/pixel) in Electron Event Representation (EER) format over 6.0 s for a total dose of 44.71 e^−^/Å^2^.

The data were processed using CryoSPARC.^24^ Briefly, the micrographs were first motion corrected using patch motion correction, followed by CTF estimation using patch CTF estimation, and particles were picked using a blob picker followed by template picking. For LarA_*Bw*_, a total of 219,896 particles were used for 3D refinement with C4 symmetry. For LarA_*Sp*_, a total of 571,110 particles were used for 3D refinement with C4 symmetry. The overall resolution was estimated based on the gold-standard Fourier shell correlation (FSC_0.143_).^25^ The final maps were deposited into the Electron Microscopy Data Bank (EMDB). The data processing procedures are described in **Figure S8** for LarA_*Bw*_ and **Figure S9** for LarA_*Sp*_. Initial models were generated using ModelAngelo in sequence mode.^26^ Refinement was carried out using Phenix,^27^ and model adjustments were carried out in Coot.^28^ Residues 4-211, 233-407, and 412-480 and residues 2-481 were modeled in the structures of LarA_*Bw*_ and LarA_*Sp*_, respectively. Model parameters were monitored using MolProbity in Phenix, and the values are listed in **Table S2** along with the respective EMD and PDB IDs. Representative density maps of both proteins are shown in **Figure S10**.

### Crystallization and structure determination of LarA_Sp_

The peak fractions from SEC were concentrated to 19 mg/ml for crystallization screening. Crystals were obtained at 21 °C with a reservoir buffer containing 0.49 M sodium phosphate monobasic monohydrate and 0.91 M potassium phosphate dibasic, pH 6.9 and flash frozen in liquid nitrogen. The diffraction data were collected at the National Synchrotron Light Source II (NSLS-II) of Brookhaven National Laboratory on the 17-ID-2 (FMX) beamline. The dataset was indexed, integrated and scaled using XDS from FastDP pipeline at NSLSII. Molecular replacement was performed to solve the structure using a monomer from the cryo-EM structure as the search model. Refinement was performed using Phenix.refine and the model was built using Coot iteratively. Crystallographic statistics are listed in **Table S3**.

## Acknowledgement

We thank Prof. Dennis W. Wolan for providing us with plasmid pET23b expressing LarA_*BW*_. We thank the MSU RTSF Cryo-EM Core Facility for data collection and processing. We thank the 17-ID-2 beamline scientist in NSLS-II for diffraction data collection and processing. This work was supported by National Institutes of Health GM128959 (to R.P.H. and J.H.), GM128959 (to R.P.H.), GM140931 (to J.H.), GM140803 (to K.N.P.) and the UCLouvain Fonds Spécial de Recherche (to B.D.).

## Author contributions

J.H., R.P.H., and K.N.P. conceived and supervised the project, and designed the experiments. S.G., D.S., J.U., and B.D. conducted biochemical and molecular biology experiments. S.S., S.G., and T.W. conducted structural biology studies and solved the structures. All authors analyzed the data and wrote the manuscript.

## Data and Material Availability

For cryo-EM structures, the density maps and corresponding atomic models have been deposited in the EMDB (EMD-72199 for LarA_*Bw*_ and EMD-72200 for LarA_*Sp*_) and PDB (9Q3J and 9Q3K), respectively. The atomic coordinates and structure factors of the crystal structure of LarA_*Sp*_ have been deposited in the PDB with the access code of 9Q2U. All data needed to evaluate the conclusions in the paper are present in the main text and/or in the Supplementary Information. Additional data related to this paper may be requested from the corresponding authors.

---

**Scheme 1.**
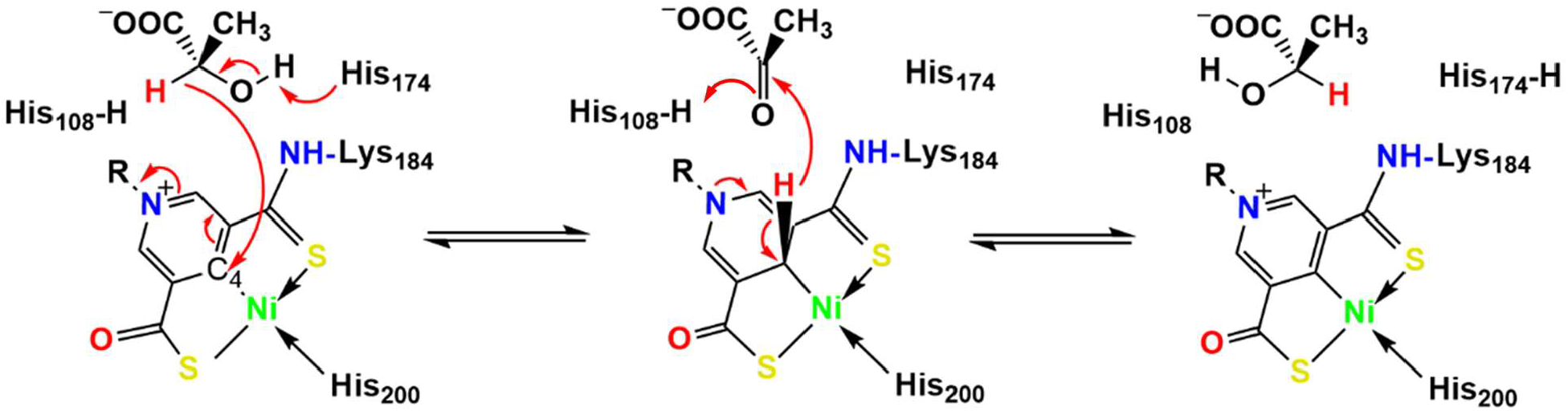
LarA_*Lp*_ catalyzed lactate racemization via a PCHT mechanism. Two catalytic residues (His108 and His174) alternately act as general acid/base. The C4 atom of the NPN cofactor acts as a hydride acceptor during lactate racemization via a pyruvate intermediate. Nickel may also transiently bind the hydride during the racemization reaction.

