## Supplementary material for "Structures of two LarA-like nickel-pincer nucleotide cofactor-utilizing enzymes with a single catalytic histidine residue": SI

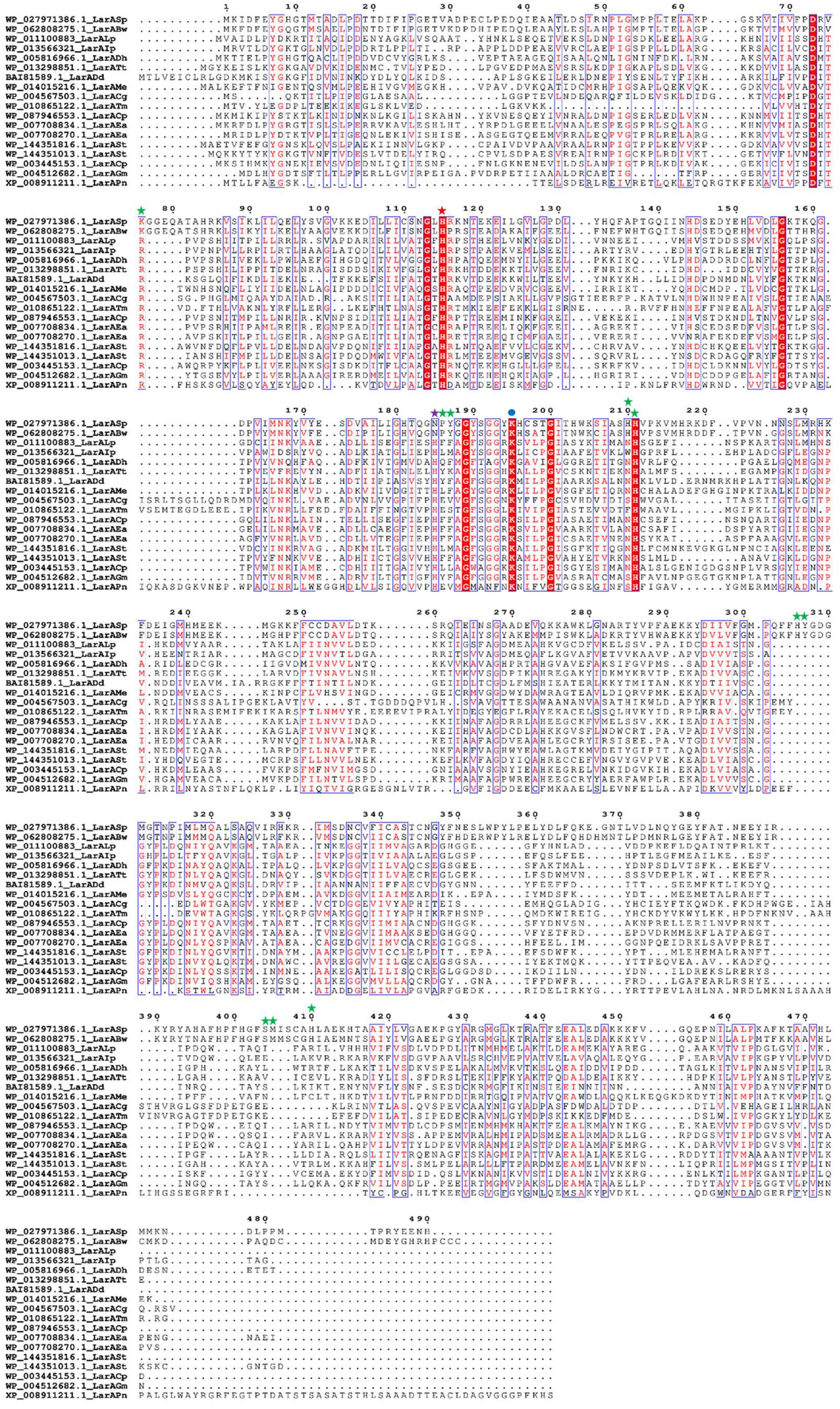

WP_027971386.1_LarASp
WP_062808275.1_LarABw
WP_011100883_LarALp
WP_013566321_LarAIp
WP_005816966.1_LarADh
WP_013298851.1_LarATt
BAI81589.1_LarADd
WP_014015216.1_LarAMe
WP_004567503.1_LarACg
WP_010865122.1_LarATm
WP_087946553.1_LarACp
WP_007708834.1_LarAEa
WP_007708270.1_LarAEa
WP_144351816.1_LarASt
WP_144351013.1_LarASt
WP_003445153.1_LarACp
WP_004512682.1_LarAGm
XP_008911211.1_LarAPn

**Figure S1.** Multiple sequence alignment of LarA$_{Bw}$, LarA$_{Sp}$, and the LarAHs with known substrates. The catalytic histidyl group and the asparaginyl side chain substituting for the second histidine residue in LarA$_{Bw}$ and LarA$_{Sp}$ are highlighted with red and purple stars, respectively. Other active site residues are indicated with green stars. The lysine residue forming the thioamide bond with the NPN cofactor in LarAHs is labeled with a blue dot. The protein IDs, species, and the primary activities are shown as follows. WP_027971386.1_LarA*Sp* from *Streptococcus plurextorum*, activity unknown; WP_062808275.1_LarA$_{Bw}$ from *Blautia wexlerae*, activity unknown; WP_011100883_LarA$_{Lp}$ from *Lactiplantibacillus plantarum*, lactate racemase; WP_013566321_LarA$_{Ip}$ from *Isosphaera pallida*, lactate racemase; WP_005816966.1_LarA$_{Dh}$ from *Desulfitobacterium hafniense*, malate racemase; WP_013298851.1_LarA$_{Tt}$ from *Thermoanaerobacterium thermosaccharolyticum*, malate racemase; BAI81589.1_LarA$_{Dd}$ from *Deferribacter desulfuricans*, hydroxyglutarate racemase; WP_014015216.1_LarA$_{Me}$ from *Megasphaera elsdenii,* hydrophobic 2-hydroxyacid racemase; WP_004567503.1_LarA$_{Cg}$ from *Corynebacterium* glutamicum, D-gluconate 2-epimerase, WP_010865122.1_LarA$_{Tm}$ from *Thermotoga maritima*, D-gluconate 2-epimerase; WP_087946553.1_LarA$_{Cp}$ from *Clostridium* pasteurianum, lactate racemase; WP_007708834.1_LarA$_{Ea}$ from *Enterocloster asparagiformis*, lactate racemase; WP_007708270.1_LarA$_{Ea}$ from *Enterocloster asparagiformis*, lactate racemase; WP_144351816.1_LarA$_{St}$ from *Sporomusa termitida*, malate racemase; WP_144351013.1_LarA$_{St}$ from *Sporomusa termitida*, hydrophobic 2-hydroxyacid racemase; WP_003445153.1_LarA$_{Cp}$ from *Clostridium pasteurianum*, hydrophobic 2-hydroxyacid racemase; WP_004512682.1_LarA$_{Gm}$ from *Geobacter metallireducens*, 2-hydroxyglutarate racemase; XP_008911211.1_LarA$_{Pn}$ from *Phytophthora nicotianae*, D-gluconate 2-epimerase.

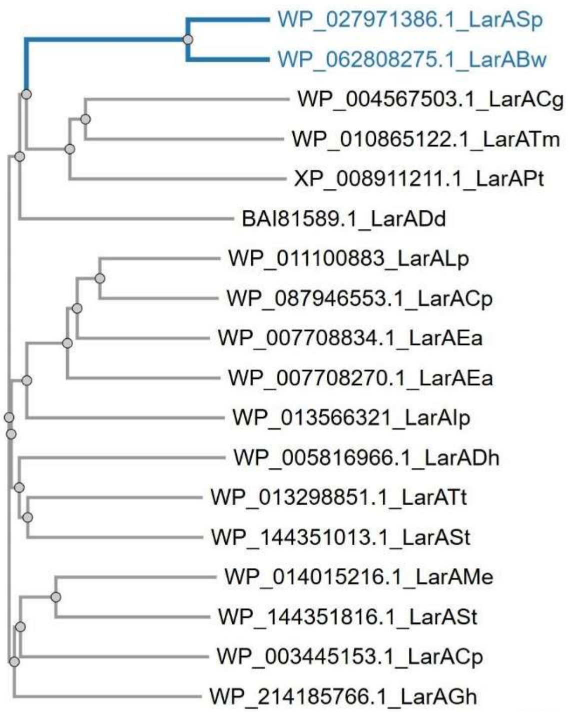


**Figure S2**. Phylogenetic analysis of LarAHs from the LarA family highlighting (in blue) the distinct branch for $LarA_{Sp}$ and $LarA_{Bw}$. The latter proteins contain an asparagine residue at the active site, whereas other LarAHs contain a catalytic histidine residue at the same position.

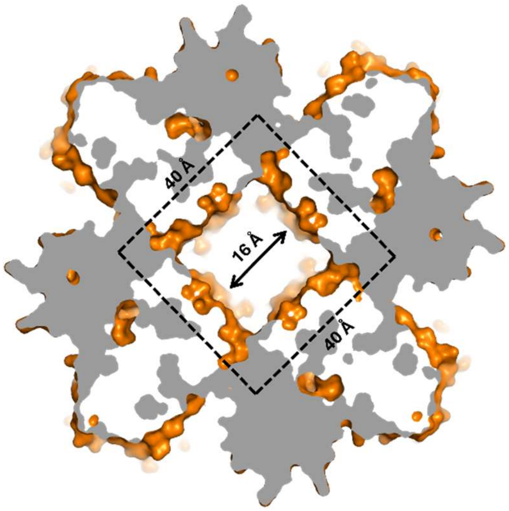


**Figure S3.** Cross-section view of the $LarA_{Bw}$ octamer in surface mode. The dashed frame indicates a square-shaped cavity with sides measuring ~40 Å within the octamer. The entrance size of the cavity is ~16 Å.

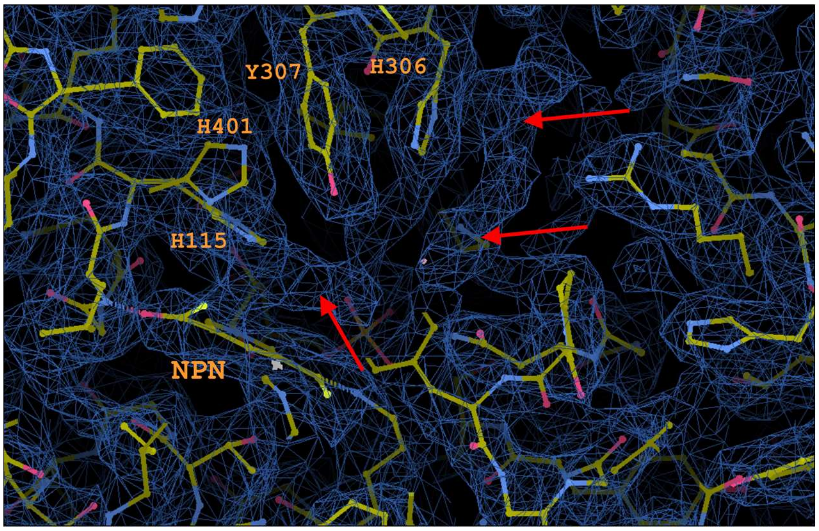


**FigureS4.** An unidentified ligand bound at the active site of LarA$_{Sp.}$ The electron densities of the ligand indicated by the arrows are close to several conserved active site residues. The contour level in Coot was set at 2.3 RMSD.

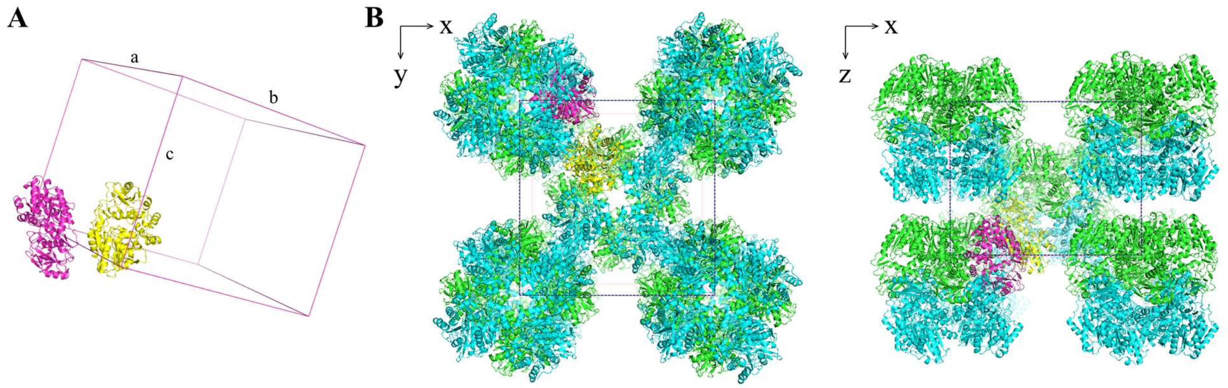


**Figure S5.** Crystal structure of LarA$_{Sp}$. (**A**) Two polypeptide chains in one asymmetric unit of a unit cell. (**B**) Octameric assembly of LarA$_{Sp}$ in the crystal lattice with a I4 space group. The rectangles indicate the unit cell boundaries.

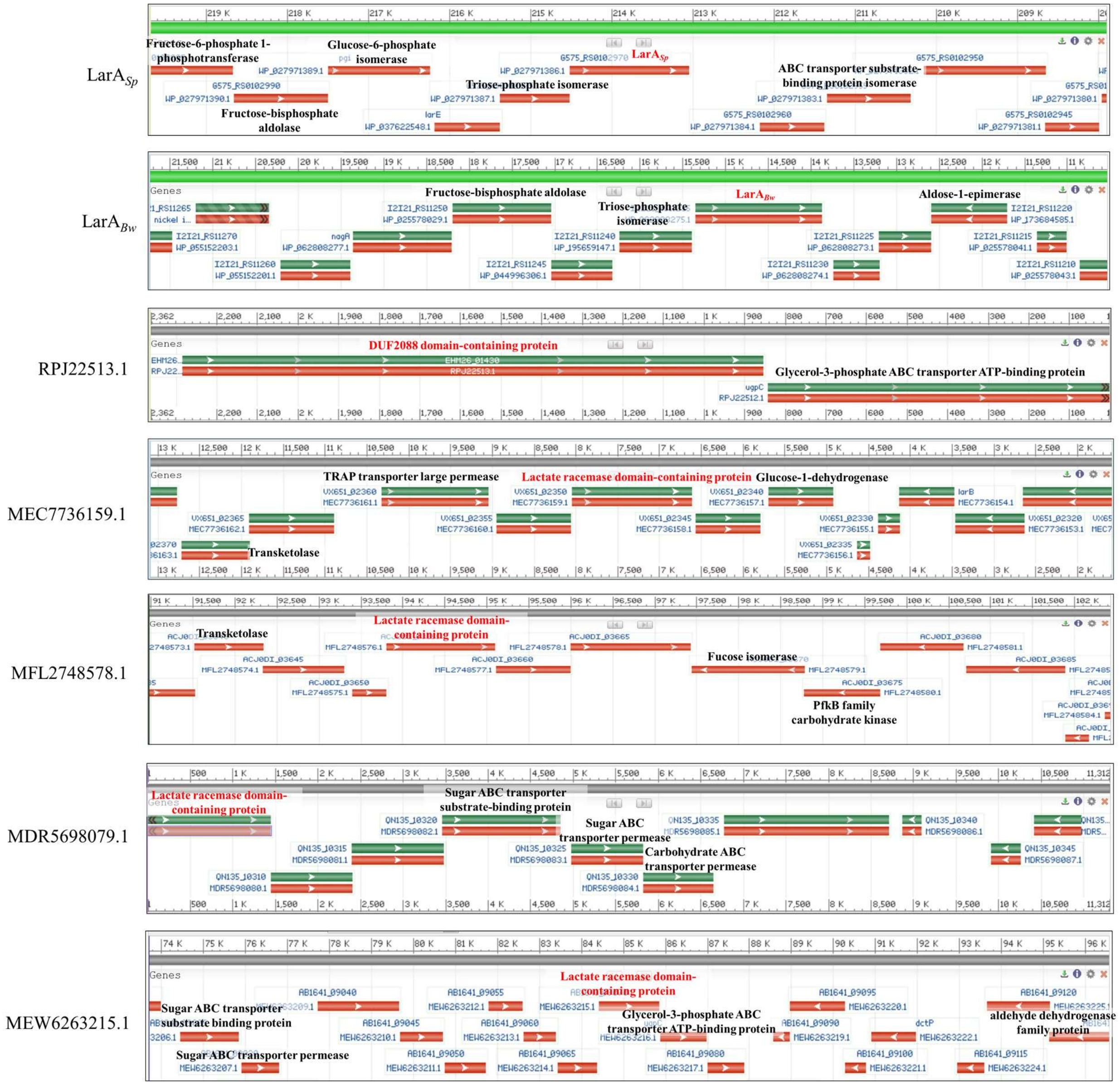


**Figure S6.** Genomic contexts of LarA$_{Sp}$, LarA$_{Bw}$ and other LarAHs from the same subfamily. These LarAHs (from **Figure 1**) share as low as 37% sequence identity. The figures were retrieved from NCBI. Only the LarAH genes (in red) and the neighboring genes annotated to be involved in carbohydrate metabolism or transport are labeled.

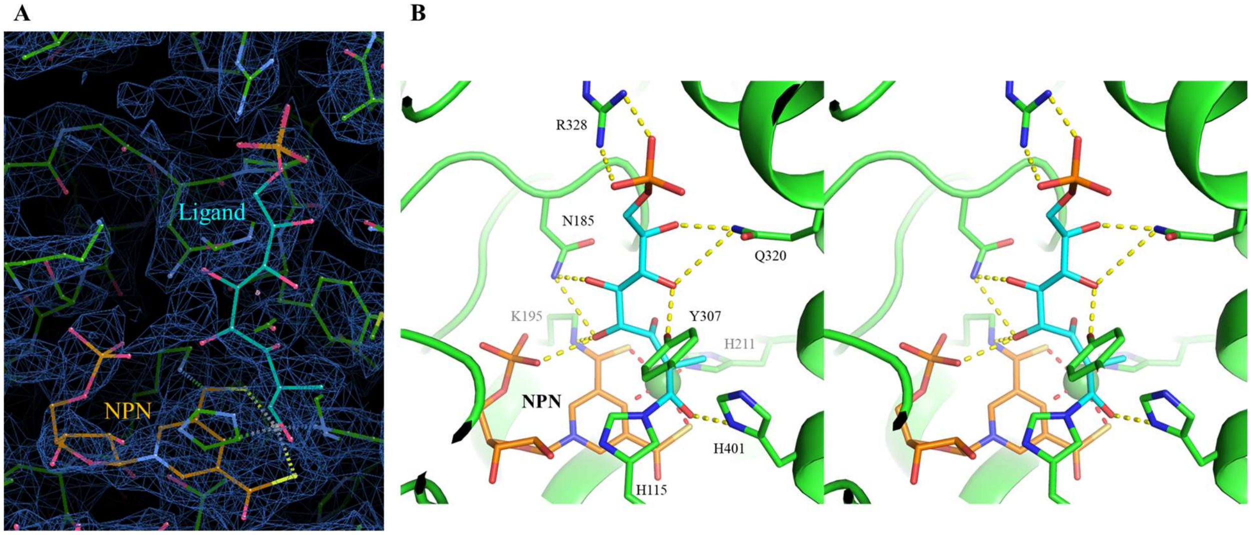


**Figure S7.** Modeling of the unidentified ligand in the active site of LarA$_{Sp}$. (**A**) Modeling of a phosphorylated aldose with a formula of $C_9H_{19}O_{10}P$. (**B**) Stereo view of the active site with the modeled ligand. The residues involved in hydrogen bond formation (yellow dashed lines) with the NPN cofactor or the ligand are labeled and shown in stick mode. The nickel ion is depicted as a green sphere. Note that this speculative modeling was used to estimate the size and shape of this ligand.

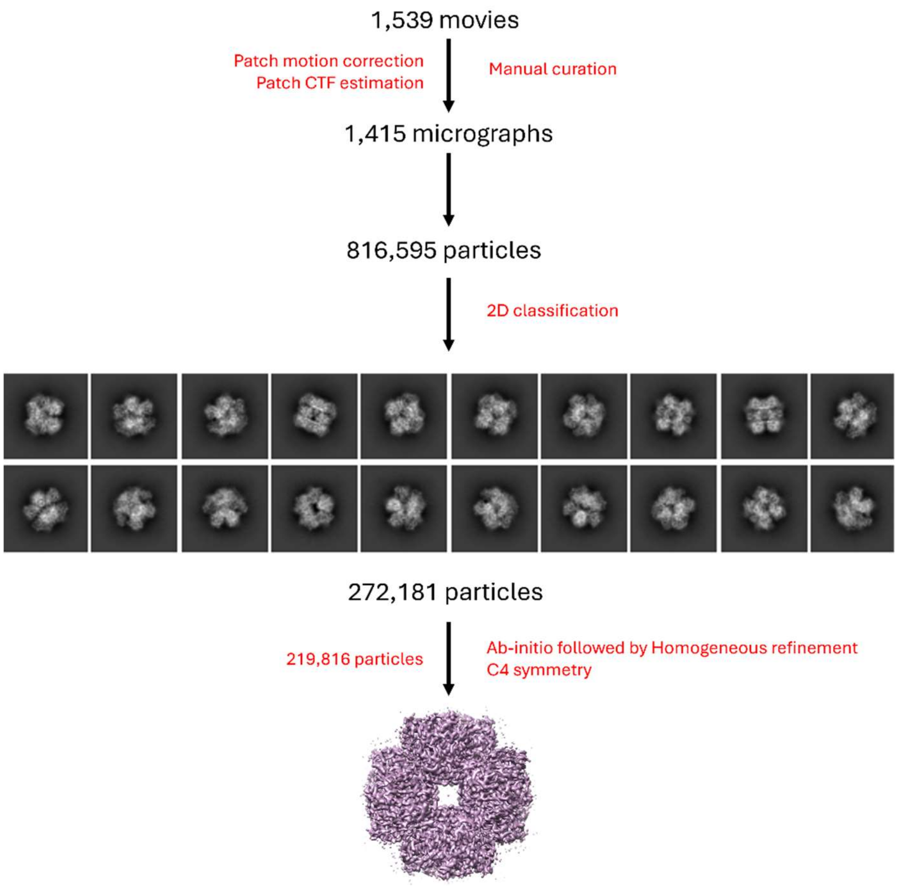


**Figure S8**. Cryo-EM data processing for $LarA_{Bw}$.

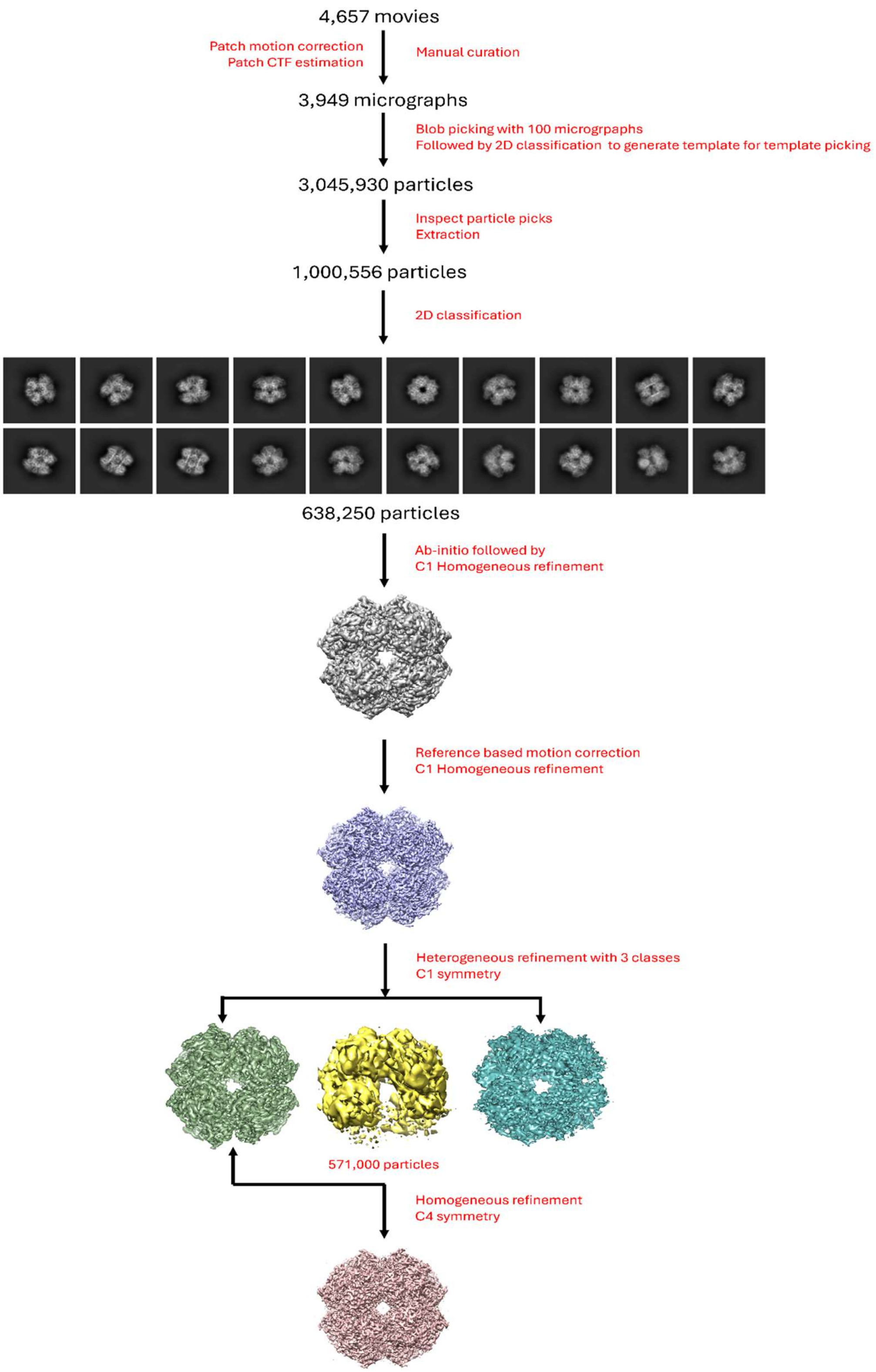


**Figure S9**. Cryo-EM data processing for $LarA_{Sp}$.

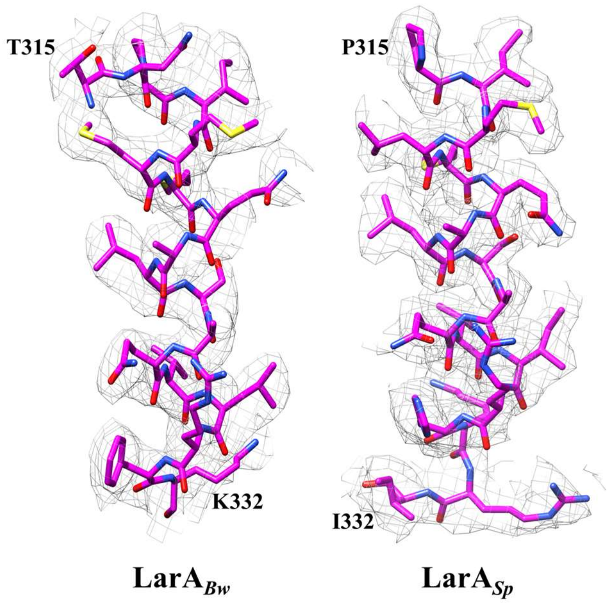


**Figure S10.** Representative cryo-EM density maps.

**Table S1.** Plasmids and primers used in this study.

| Strains, plasmids, or primers | Characteristic(s) or sequence | Source or reference |
|---|---|---|
| Strains | | |
| *L. Lactis* NZ3900 | MG1363 derivative | [1] |
| *E. coli* BL21-Gold(DE3) | | Invitrogen |
| Plasmids | | |
| pGIR213 | $Cm^r$; pGIR210 with DNA encoding $LarA_{Sp}$ | This study |
| pET23b_$LarA_{Bw}$ | $Amp^r$; pET23b with DNA encoding $LarA_{Bw}$ | [2] |
| pGIR112_PciI | $Cm^r$; pGIR112 with an introduced PciI site | This study |
| pGIR112_$LarA_{Bw}$ | $Cm^r$; pGIR112_PciI with DNA encoding $LarA_{Bw}$ | This study |
| Primers | | |
| LarAH37_A | AAAACATGTCTAAAATTGATTTTGAATACGGTCATGGG | This study |
| LarAH37_B2 | AAATCTAGAATGATTCTCCTCATAACGAGGCGTCATAG | This study |
| PciI_F | ATAAATTATAAGGAGGCACTCAACATGTCCGTTGCAAT TGATTTACCATATG | This study |
| PciI_R | CATATGGTAAATCAATTGCAACGGACATGTTGAGTGC CTCCTTATAATTTAT | This study |
| LarAH13_F | CATGCACATGTCCAGCAAATTTGATTTTGAGTATGGA C | This study |
| LarAH13_R | GATC GCT AGC ACAGCAACATGGATGACGATGTCC | This study |

**Table S2.** Cryo-EM data collection, refinement and validation statistics.

| **Map** | **LarA*Bw*** | **LarA*Sp*** |
|---|---|---|
| EMD Identifier | EMD-72199 | EMD-72200 |
| PDB Identifier | 9Q3J | 9Q3K |
| **Data collection and processing** | | |
| Magnification | 120,000 | 130,000 |
| Voltage (kV) | 200 | 200 |
| Electron exposure ($e^{-}$/Å$^2$) | 32.27 | 44.71 |
| Defocus range (μm) | -0.8 to - 3.0 | -0.8 to - 3.0 |
| Pixel size (Å) | 0.872 | 0.886 |
| Symmetry imposed | C4 | C4 |
| Number of micrographs | 1,539 | 4,657 |
| Number of particles | 219,896 | 571,110 |
| Map resolution (Å) | 3.15 | 2.20 |
| FSC threshold | 0.143 | 0.143 |
| **Refinement** | | |
| *Model composition* | | |
| Protein atoms | 28,825 | 30,325 |
| $H_2O$ molecules | 0 | 0 |
| Ni atoms | 0 | 8 |
| 4EY: P2TMN | 0 | 8 |
| *B-factors (Å$^2$)* | | |
| Overall | 35.05 | 45.30 |
| Protein atoms | 35.05 | 45.25 |
| $H_2O$ molecules | 0 | 0 |
| Ni atoms | 0 | 86.09 |
| 4EY: P2TMN | 0 | 51.20 |
| *R.m.s. deviations* | | |
| Bond lenghts (Å) | 0.003 | 0.003 |
| Bond angles (°) | 0.557 | 0.574 |
| *Validation* | | |
| MolProbity score | 1.57 | 1.75 |
| Clashscore | 6.77 | 17.23 |
| Poor rotamers (%) | 0.16 | 0.97 |
| *Ramachandran* | | |
| Favored (%) | 96.83 | 97.93 |
| Allowed (%) | 3.17 | 2.07 |
| Outliers (%) | 0.00 | 0.00 |
| CC (volume) | 0.85 | 0.82 |

**Table S3.** Crystallographic statistics for $LarA_{Sp}$.

| | |
|---|---|
| **Data collection** | |
| Beamline | NSLSII 17-ID-2 FMX |
| Wavelength (Å) | 0.97934 |
| Space group | I4 |
| Unit cell a, b, c (Å);<br>α, β, γ (°) | 145.224, 145.224,<br>116.994<br>90.00, 90.00, 90.00 |
| [a]Resolution (Å) | 34.23 – 3.108<br>(3.16-3.108) |
| [a]Redundancy | 14.1 (14.8) |
| [a]Completeness (%) | 100 (100) |
| [a]$I/\sigma I$ | 7.3 (0.7) |
| [a,b]$R_{merge}$ | 0.326 (4.635) |
| [a,c]$R_{pim}$ | 0.090 (1.244) |
| [d]$CC_{1/2}$ | 0.995 (0.364) |
| **Refinement** | |
| Unique reflections | 21,937 |
| Number of atoms | 7183 |
| Protein atoms | 7162 |
| $H_2O$ molecules | 21 |
| [e]$R_{work}/R_{free}$ | 0.222/0.282 |
| *B*-factors (Å$^2$) | 94.6 |
| Protein atoms | 94.7 |
| $H_2O$ molecules | 72.9 |
| R.m.s. deviation in bond lengths (Å) | 0.0124 |
| R.m.s. deviation in bond angles (°) | 1.47 |
| Ramachandran plot (%) favored | 90.10 |
| Ramachandran plot (%) allowed | 9.80 |
| Ramachandran plot (%) outliers | 0.10 |
| Rotamer (%) outliers | 0 |
| PDB ID | 9Q2U |

[a]Highest resolution shell is shown in parentheses.
[b]$R_{merge} = \sum_{hkl}\sum_j |I_j(hkl)-\langle I(hkl)\rangle| / \sum_{hkl}\sum_j I_j(hkl)$, where $I$ is the intensity of reflection.
[c]$R_{pim}=\sum_{hkl} [1/(N-1)]^{1/2}\sum_j |I_j(hkl)-\langle I(hkl)\rangle| / \sum_{hkl}\sum_j I_j(hkl)$, where N is the redundancy of the dataset.
[d]$CC_{1/2}$ is the correlation coefficient of the half datasets.
[e]$R_{work} = \sum_{hkl} |\,|F_{obs}| - |F_{calc}|\,| / \sum_{hkl} |F_{obs}|$, where $F_{obs}$ and $F_{calc}$ is the observed and the calculated structure factor, respectively. $R_{free}$ is the cross-validation R factor for the test set of reflections (5% of the total) omitted in model refinement.
